# Sensory prediction error drives subconscious motor learning outside of the laboratory

**DOI:** 10.1101/2023.02.22.529399

**Authors:** Scott T. Albert, Emily Blaum, Daniel H. Blustein

## Abstract

Sensorimotor adaptation is supported by at least two parallel learning systems: an intentionally controlled explicit strategy, and an involuntary implicit learning system. To investigate the error sources driving these two systems, past work focused on constrained reaches or finger movements in laboratory environments has shown subconscious learning systems to be driven in part by sensory prediction error (SPE), i.e., the mismatch between the realized and expected outcome of an action. We designed a ball rolling task to explore whether SPEs can drive implicit motor adaptation during complex whole-body movements that impart physical motion on external objects. After applying a visual shift, participants rapidly adapted their rolling angles to reduce the error between the ball and target. We removed all visual feedback and told participants to aim their throw directly toward the primary target, revealing an unintentional 5.06° implicit adjustment to reach angles that decayed over time. To determine whether this implicit adaptation was driven by SPE, we gave participants a second aiming target that would ‘solve’ the visual shift, as in Mazzoni and Krakauer (2006). Remarkably, after rapidly reducing ball rolling error to zero (due to enhancements in strategic aiming), the additional aiming target caused rolling angles to deviate beyond the primary target by 3.15°. This involuntary overcompensation, which worsened task performance, is a hallmark of SPE-driven implicit learning. These results show that SPE-driven implicit processes, previously observed within simplified finger or planar reaching movements, actively contribute to motor adaptation in more complex naturalistic skill-based tasks.

**New and Noteworthy:** Implicit and explicit learning systems have been detected using simple, constrained movements inside the laboratory. How these systems impact movements during complex whole-body, skill-based tasks has not been established. Here we demonstrate that sensory prediction errors significantly impact how a person updates their movements, replicating findings from the laboratory in an unconstrained ball-rolling task. This real-world validation is an important step towards explaining how subconscious learning helps humans execute common motor skills in dynamic environments.

## Introduction

Current theories posit that implicit, i.e., subconscious, motor learning is driven in part by sensory prediction error (SPE): a mismatch between a movement’s outcome versus the expected result (1– 7). The notion that errors are a critical implicit learning substrate is central to both descriptive and computational motor control models (1, 3, 8–10). The visuomotor rotation (VMR) paradigm has been applied extensively to examine error-based adaptation in simplified motor tasks where participants perform a planar reaching or pointing movement to advance a cursor towards a virtual target (Fig 1, Inset). For example, SPE-driven learning was initially detected by Mazzoni and Krakauer (2), in an experimental condition where the arm was splinted to a tripod, and the movement of a single index finger was tracked. While these constraints on task complexity allow precise error manipulation (2, 7), the SPE-learning mechanisms detected in these contexts may not generalize to skill learning in natural environments, where individuals must coordinate multiple body segments to impart a desired motion on an external object. In other words, do SPEs contribute to our motor control and learning within full-body actions we perform in day-to-day life, like soccer or bowling?

**Figure 1.**
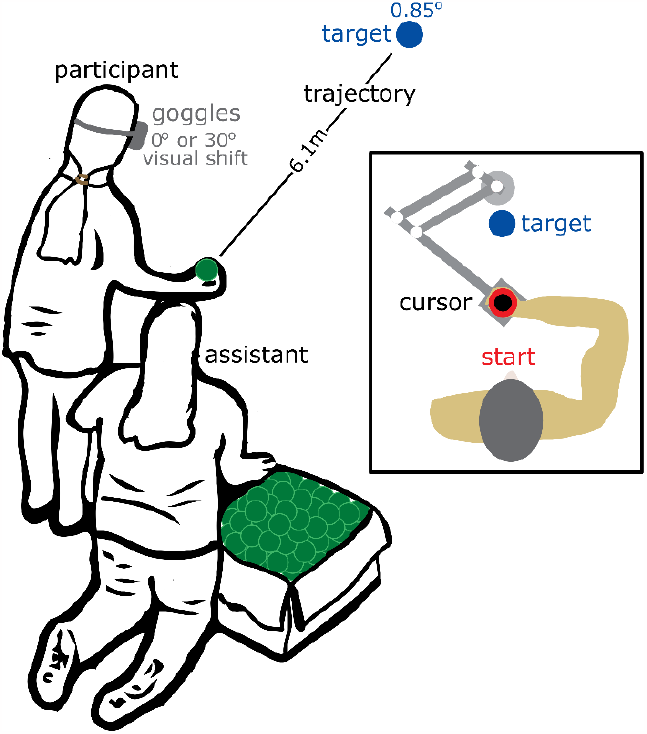
Unconstrained motor task setup. Participants wearing goggles with or without a visual shift rolled a tennis ball to a small target 6.1 m away on sequential trials. **Inset**. Traditional VMR experiment (adapted from (48)).

In the rare cases where sensorimotor adaptation has been examined in skill-based settings (11–14), reliable assays to separate implicit learning from explicit (i.e., intentional) strategies have not been applied; these studies have either relied on the tenuous assumption that implicit learning alone persists once a perturbation has been removed, or have attempted to reduce explicit learning by providing aiming instructions, an intervention that could cause the opposite effect: priming explicit adjustments (13). Furthermore, no attempts have been made to dissociate between error sources that could drive the implicit system, in particular, SPEs versus target (i.e., performance) errors (1, 3, 7, 10, 15) which co-occur unless specialized conditions are used to isolate the SPE component.

Here we aimed to fill this gap using a task designed to detect SPE learning in a more naturalistic skill-based task that requires complex multi-segmented body movements. By eliminating target errors and isolating SPEs, we observed an obligatory overcompensation in motor actions, demonstrating that subconscious adjustments to everyday movements are driven at least in part by SPEs.

## Methods

### Ethics statement

Human participant research was completed with the approval and oversight of the Rhodes College Institutional Review Board.

### Subject recruitment

Thirty study participants were recruited by word of mouth, and each given a $10 gift card for their participation (mean age = 20yrs; age range – 18 – 21; 21 female; 29 right handed but all completed ball throws with right). Data from three participants were omitted because of technical issues that resulted in missing video files that contained motor error data. Written informed consent was obtained from each participant before completing the experiment.

### Experimental setup

Participants stood behind a line marked on the floor facing a small target (D=0.85°) on the ground 6.1m away. They were informed about the task parameters: to use an underhand motion to roll a tennis ball so that it moved through the target. The experiment took place in a black box theater with black flooring and a black curtain behind the target area under which balls rolled out of view. Participants started by wearing goggles with clear lenses that allowed for unperturbed vision (Fork in the Road Vision Rehabilitation Services LLC, Madison, WI). During shifted vision blocks, participants wore Perception (Distortion) goggles (WinginItProducts, Pleasant Prairie, WI) that caused the participants to perceive visual targets offset in space by 30° from their true position. Both types of goggles occluded the view of the body and throwing arm and participants were asked to release the tennis ball below their waist level so the hand was never visible.

An infrared video camera (Prime Color, Optitrack) was mounted directly above the target (∼7 meters above attached to a theater lighting grid). The camera was pointed at the target, capturing several meters of linear space on both sides of the target. It recorded the tennis ball rolling along the floor so that rolling error could be computed (see Data analysis). The camera was attached to a computer which displayed and recorded the video in Motive software (Optitrack). A single eStrobe (Optitrack) mounted above the target (next to the camera) provided the only illumination in the room and could be toggled on and off with a single button press in the Motive software. The camera system did not record any motion tracking of participant kinematics.

A researcher kneeled next to the participant and handed tennis balls to the participant for each subsequent roll. Another researcher stood behind the curtain and collected tennis balls to make sure they did not roll back onto the visible floor space. A third researcher monitored the video recording and controlled the lights during no feedback trials. For some participants, only two researchers were present and the tennis ball wrangling and computer monitoring were done by the same individual.

*Data collection*. Each participant completed two blocks of 192 tennis ball rolls. In the ‘No Instructions’ block, participants were not given a strategy to counteract the visual perturbation. In the ‘Aiming Target’ block, participants were given a secondary target to use that would minimize the error resulting from the visual perturbation. The order of the blocks was randomly assigned to the participants (after the three excluded participants, 13 participants completed ‘No Instructions’ first and 14 participants completed ‘Aiming Target’ first).

Participants first completed 40 baseline movement trials, rolling the ball to the target without shifted vision. Then participants switched to goggles that shifted their vision by 30°. To switch the goggles, participants first closed their eyes. The researcher then assisted with doffing the current goggles and donning the new goggles. With eyes still closed, participants were then guided through a series of in-place rotations in order to reset the relative orientation between the target and the body. Then participants opened their eyes.

In the ‘No Instructions’ block, participants then completed two trials with shifted vision after which the researcher said: “Try to hit the target.”. Participants then completed another 70 trials with the shifted visual feedback.

In the ‘Aiming Target’ block, participants completed two trials with shifted vision after which the researcher placed an aiming target, matching the size and appearance of the original target, 30° to the right of the original target. The researcher then explained to the participant: “You just made two large errors because we imposed a rotation that pushes you 30 degrees to the left. You can reduce this error by aiming at the new target to your right.”

After 72 total trials with the shifted goggles, participants then completed 10 trials with the same visual shift but with no endpoint error feedback. Participants were made aware before these trials that the light would turn off immediately after they released the ball. The researcher controlling the computer manually shut off the eStrobe light as soon as the tennis ball first hit the floor at the start of the participant’s roll, making the room completely dark. Since the camera was imaging in the infrared spectrum, the tennis ball could still be viewed from the computer. As soon as the tennis ball passed the target and went out of view underneath the curtain, the researcher turned on the light.

After 10 trials with no endpoint visual feedback, the goggles were switched back to the clear lens goggles and participants were told: “I would like you to aim for the original target again.” Participants completed 10 additional trials without endpoint visual feedback with no vision shift. Then participants completed 60 trials with no visual shift with endpoint visual feedback (i.e. the lights stayed on).

### Data analysis

Video recordings were analyzed using Vernier Video Analysis software (Vernier Science Education, Beaverton, OR). Small pieces of tape on the ground set at 1m apart and visible in the video frame but not visible to participants were used to calibrate position measurements in the software. When the tennis ball was in line with the target, its position was marked manually with a mouse click in the software for each trial. The lateral error between the target and tennis ball was computed for the entire set of trials using the marked positions on the video frames.

Data were loaded into MATLAB software and a custom script was used to calculate descriptive statistics and identify the steady-state trials during the visual rotation block after which learning had stabilized (16). Total Adaptation was calculated as the sum of the visual shift (30°) and the mean error over the last three trials of the shifted vision with feedback trials [i.e. how much movements were adapted under the visual shift]. Implicit adaptation was calculated as the mean error over the first three trials immediately after switching to unshifted vision (after the shifted vision block). Explicit adaptation was computed as the difference between Total Adaptation and Implicit Adaptation. Graphical plots were generated in R and MATLAB supported by publicly available scripts (17, 18).

A multiple linear regression was run in RStudio (v2022.07.1, R version 4.1.1) with Total Adaptation and Implicit Adaptation as the outcome variables and Block Number and Strategy Condition (i.e. with or without aiming target) as the predictor variables. Explicit Adaptation was not included in the regression since it is computed from the included outcome variables and thus correlated with them. Secondary inferential statistical analyses were conducted to further explain the results. Bonferroni-corrected p-values from t-tests were used to describe pairwise comparisons that appear in Fig. 2C,D. The same analysis was run on pairwise comparisons with data separated into the four different blocks (Fig. 3). The mean tennis ball error during the last three trials of the rotated vision trials with feedback was analyzed using a one sample t-test to determine if the ball’s error was beyond the physical extent of the target (i.e. including the radius of the target and ball). To investigate savings (and by extension, the influence of an explicit strategy), a paired t-test was run comparing initial error on the first two visual shift trials between Block 1 and Block 2. To investigate individual correlations between total and implicit adaptation rates, Pearson’s correlation coefficients were computed for no-aiming-target and aiming-target data.

**Figure 2.**
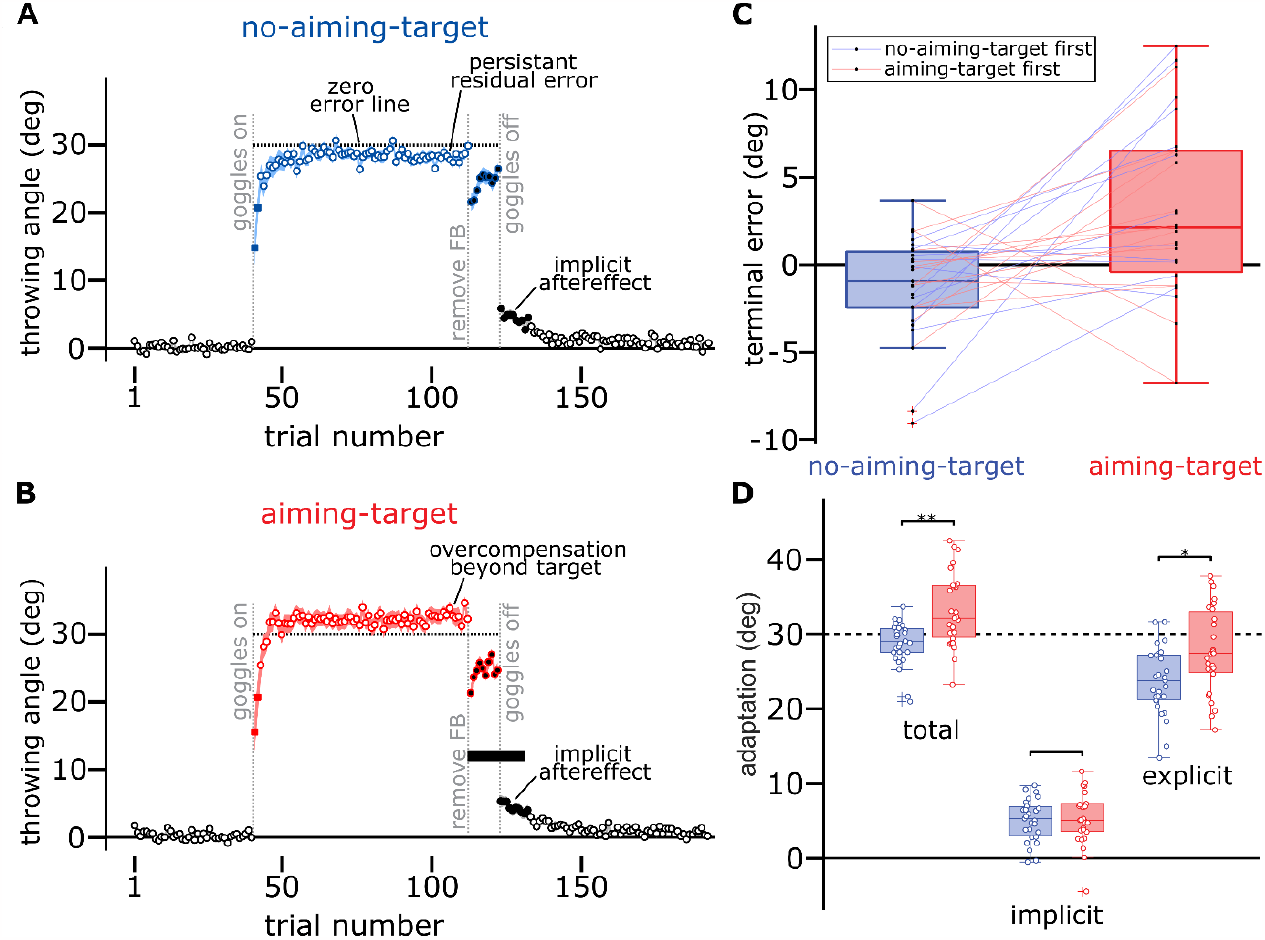
Implicit and explicit components of unconstrained movements. **A**. Trial-by-trial lateral error during ball rolling relative to the perceived target location with no aiming target provided (n = 27). Black-outlined datapoints represent trials with no visual shift applied. Blue-outlined points represent trials with a 30° visual shift applied. After the first two visually shifted trials (blue squares), participants were reminded of the goal to hit the target. Black-filled datapoints indicate the absence of endpoint visual error feedback. **B**. Trial-by-trial error for participants given an aiming target to minimize error (n = 27). Red-outlined datapoints indicate trials with a 30° visual shift applied. After the first two visually shifted trials (red squares), the aiming target was added and described. **C**. Mean residual error on the last three trials of the visual shift block deviated beyond the target for the aiming-target group (red boxplot). Color-coded lines connect each individual participant’s results for both conditions: blue lines represent participants who completed the no-aiming-target condition first; red lines represent participants who completed the aiming-target condition first. **D**. Statistically significant differences between aiming-target and no-aiming-target groups in total adaptation and explicit adaptation. Error bars represent standard error and asterisks indicate results of Bonferroni-corrected p-values from paired t-tests [* p=0.012; ** p=0.0005].

**Figure 3.**
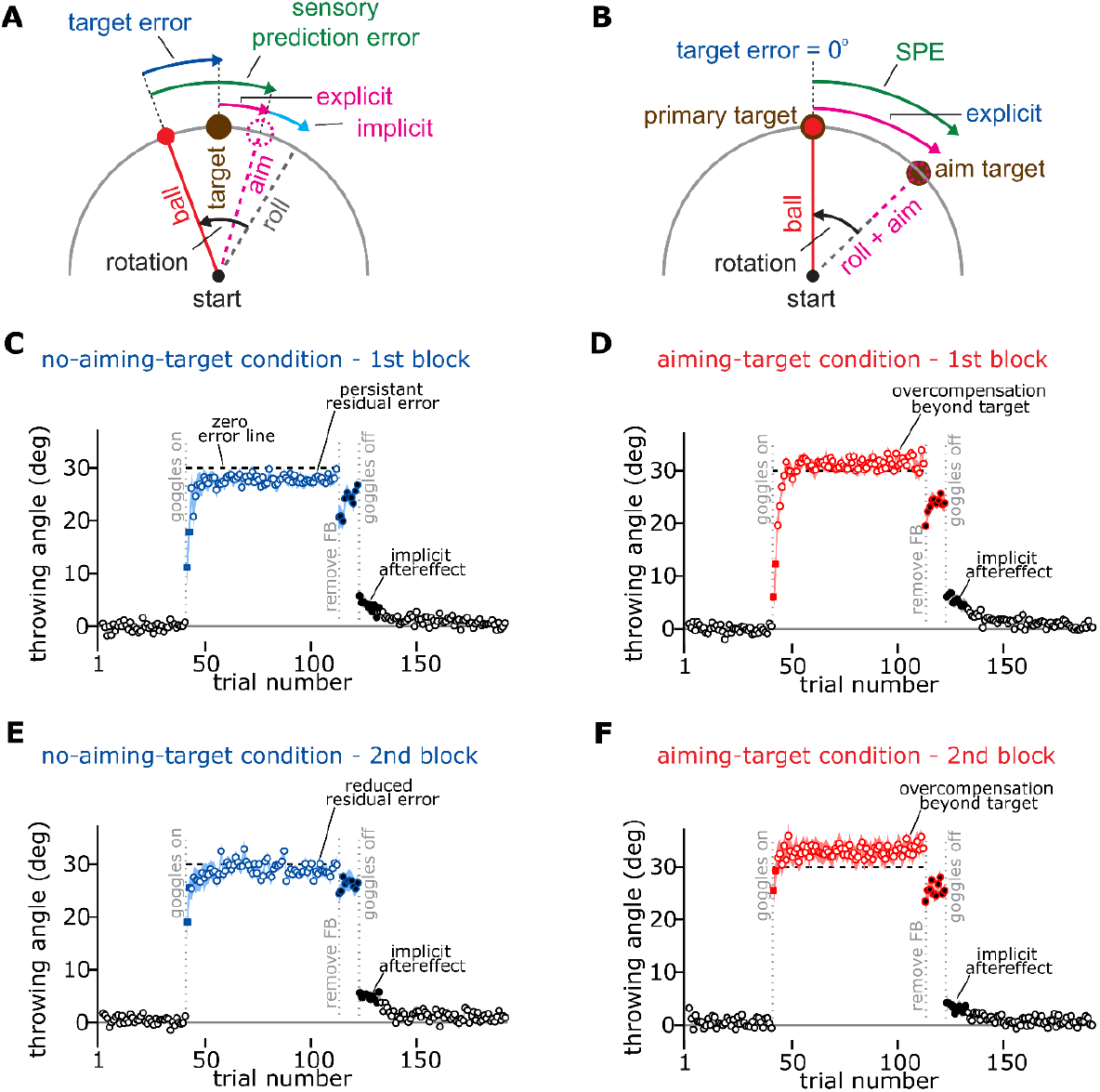
Trial-by-trial error across all block order and strategy condition pairs. **A**. Overview of error and adaptation components for the no-aiming-target condition. **B**. Overview of error and adaptation components for the aiming-target condition. **C**. Data plotted for participants completing their first block without an aiming target (n = 13). Plotting conventions as described in Figure 2. **D**. Data from Block 1 of participants using an aiming target (n = 14). **E**. Data from Block 2 of participants without an aiming target who first used an aiming target, the same participants as in panel B (n = 14). **F**. Data from Block 2 participants with an aiming target who first had no aiming target, the same participants as in panel A (n = 13).

### Data and code

All data, analysis code and plotting scripts are available here: https://osf.io/8kf59/?view_only=cfeb2f109d304994bbd3a5f5a44c11ae

## Results

We trained participants to roll a tennis ball (D = 0.425° at target displacement) towards a small target (D=0.85°) 6.1m away; the ball arrived at the target about 750ms after release (Fig. 1). Unlike constrained motor tasks where motion is restricted to one or two joints (2, 7, 19), our paradigm introduced an array of additional control variables: wrist, elbow, and shoulder angle, bend at the knees and hip, orientation of the trunk, release timing, grip, and arm speed. Further, to internalize sensory error, the brain needed to map visual error between the ball and target onto the participant’s temporally separated motion (conditions known to attenuate implicit learning (8, 20, 21)).

Despite these complex elements, participants achieved excellent performance, hitting the target with high precision (Fig. 2A,B, first 40 trials; 0.095° average error [mean across all 27 participants of median error on last 10 trials of initial baseline trials]). Next, we rotated the participant’s visual scene with prism glasses. The glasses, which occluded the view of the body and throwing arm, caused participants to perceive the target offset by 30° in space from its true position. Past the target, balls disappeared underneath a black curtain, which along with black flooring, reduced spatial reference cues. In response, participants committed throwing errors (Fig. 2A, blue rotation period) which induced rapid adaptation; participants adjusted their movement angle to mitigate the discrepancy between ball and target, reaching asymptotic performance (16) by approximately their 13^th^ trial with -1.29° error persisting over the last 3 trials of the period (Fig. 2A,C).

Was this rapid motor adaptation solely dependent on explicit changes in throwing strategies, e.g., intended aiming angle, or was some adaptation due to subconscious implicit learning? To measure implicit and explicit learning components, we instructed participants to continue making throws over 20 trials, but without error feedback. On these trials, participants could view the initial target location, but the room was darkened once the ball left the hand. For the initial 10 no-feedback trials, the rotation was maintained. Then, participants switched to the unshifted glasses for an additional 10 no-feedback trials. For these trials, participants were told that the glasses had been switched, and that they should aim towards the initial target location. We gave these instructions to match the conditions used to measure implicit learning via aftereffect trials in modern VMR experiments (1, 8, 22). Throwing angles decreased from 28.71° (relative to perceived target location) at the end of the adaptation period, to approximately 5.06° on the initial 3 unshifted no-feedback trials (Fig. 2A, implicit aftereffect across Trials 11-13 of the no-feedback period; Trials 1-10 were not used as vision remained shifted and no aiming instruction had been provided). This immediate voluntary change in angle implied that participants had explicitly aimed about 23.7° away from the perceived visual target (explicit strategy was estimated by subtracting implicit learning from total adaptation during the shifted vision block). An additional 5° of subconscious implicit learning decayed after a return to unshifted vision, both in the absence of feedback (one sample Bonferroni-corrected t-test confirming the slope of error over these trials to be significantly different from zero, t(26) = -1.54, p= 0.0097) and over the ensuing washout period (t(26)= -2.74, p < 0.0001).

In sum, motor behavior in our task showed a clear correspondence to lab-based VMR studies: 1) Participants adapted to the rotation, achieving steady-state performance that never completely counteracted the rotation (3, 8, 23). 2) Learning was supported by two parallel systems (2, 7, 24, 25): an explicit strategy that participants could willfully disengage, and an implicit correction that persisted despite the participant’s conscious intentions. But which error sources had engaged the implicit learning system? Implicit learning could have been caused by SPEs (error between predicted and realized motion), task errors (error between ball and target) (1, 3, 10, 15), or a model-free reinforcement of successful motor actions (26).

While explicit strategies are driven by errors in performance (interchangeably called task, target, or performance error) (3, 10, 25, 27-29) implicit learning is modulated by both task errors (Fig. 3A; error between ball and target) and SPEs (Fig. 3A; error between the ball and aiming vector) (1, 2, 3, 7, 10, 15). Generally, these error sources cannot be easily disentangled because they co-occur and are oriented in the same direction (note how both SPE and target error point in the same direction in Fig. 3A). This issue, however, can be resolved by eliminating target error: a condition achieved in past studies (2, 7) by providing a second aiming target that precisely counteracts the rotation (see Fig. 3B; aiming to the “aim target” reduces the target error to 0°, leaving only SPE).

Thus, to test whether SPEs contributed to adaptation we tested subjects in a second paradigm similar to Mazzoni and Krakauer (2006) (Fig. 2B). Here, after performing two ball rolls with rotated vision, participants were told they could aim towards an assistive target in order to move the ball to the desired goal target. Subjects in this aiming-target condition adopted this strategy quickly, reaching steady-state performance in about 10 trials (see Methods (16)). Remarkably, however, despite rapidly achieving zero error (i.e., hitting the target’s center), the ball angle rapidly surpassed the target’s location (horizontal dashed line in Fig. 2B), overcompensating for the imposed rotation. By the last 3 trials of the period, the ball’s path deviated by 3.15° *beyond* the task-related target (Fig. 2C), exceeding on average the target’s extent (radius = 0.425°, t(26) = 2.89, p = 0.0077, 95% CI lower bound = 1.21°). This involuntary overcompensation in reach angle is a distinct hallmark of SPE-driven implicit learning, indicating that SPE learning mechanisms detected in simplified arm and hand movements studied in the laboratory, also contribute to subconscious adaptation in skill-based tasks involving multi-segmented full-body motion and object manipulation.

To examine why this involuntary overcompensation occurred only when a secondary aiming target was provided, we compared implicit and explicit learning measures across the aiming-target and no-aiming-target conditions. As expected, the increase in total adaptation observed in the aiming-target condition (main effect of aiming target on total adaptation with a multiple linear regression; F(2,51) = 10.00, p = 0.00022 for total adaptation response variable; β_aiming-target_ = 4.51, t(24) = 4.19, p = 0.00011), was supported by an increase in explicit strategy (Fig. 2D; no aiming target: 23.64°; aiming target: 27.85°; t(52) = -3.00, p = 0.0123; follow up paired t-test with Bonferroni correction, see Methods); thus, participants utilized the aiming target and instructions to increase their explicit strategy. On the contrary, the aftereffects observed immediately upon returning to unshifted vision, indicating implicit adaptation, did not show a statistically significant difference between groups (5.06° (no aiming target), 5.30° (aiming target) (F(2,51) = 0.89, p = 0.42 for implicit adaptation response variable). This indicated that the SPEs related to the goal target (no-aiming-group) and aiming-target (aiming-group) both drove implicit learning in a similar manner, suggesting that implicit learning was largely immune to any change in explicit strategy in our experiment. This was further supported at the individual participant level; we did not observe that individuals with greater implicit aftereffects exhibited greater total adaptation, and hence overcompensation. Specifically, we observed only weak correlations between total adaptation and implicit adaptation across the no-aiming-target condition (r(25) = -0.25, p = 0.20) and the aiming-target condition (r(25) = 0.098, p = 0.63). In sum, overcompensation appeared to be due to an overactive explicit process that did not appropriately counterbalance obligatory SPE-driven learning in the implicit learning system.

This overcompensation pattern appeared largely insensitive to the order in which each condition was experienced (i.e., all participants completed the aiming-target and no-aiming-target periods, but with their order counterbalanced). We did not observe a statistically significant increase in overcompensation when aiming targets were provided during the second learning period (4.31°, Fig. 3F) than the first learning period (2.06°, Fig. 3D) (t-test: t(25) = -1.20, p = 0.24). Similarly, during the no-aiming-target condition, we also did not detect a statistically significant change in residual error between the first (Fig. 3C, -2.05°) and second (Fig. 3E, -0.59°) learning periods (independent t-test: t(25) = -1.33, p = 0.20). On the contrary, initial learning rates were highly sensitive to the block order; both the aiming-target and no-aiming target conditions exhibited an increased learning rate when these periods were experienced during the second block of the experiment, likely owing to a savings in explicit strategy (30–32) upon re-exposure to the rotation [mean error on the first two shifted vision trials on Block 1 (−19.36°, Fig. 3C,D) exceeded that in Block 2 (−4.71°, Fig. 3E,F); paired t-test (t(26)=-8.09, p<0.0001]. These increases in overall adaptation during the early phase of adaptation, but not the later steady-state period suggests that participants downregulated strategic aiming over time, likely to mitigate the obligatory SPE-driven increases in the implicit process (24).

## Discussion

Error-based motor adaptation has been studied extensively in laboratory experiments that constrain motor tasks to the planar motion of a finger, hand, or forelimb. Little is known about whether this vast body of work applies to more complex skill-based motor actions we perform in day-to-day life involving coordination across multiple segments across our entire body. Here, we observed the first direct evidence that 1) subconscious error-driven learning processes make substantial contributions to more sophisticated motor actions, and 2) that this learning is in part mediated by SPEs not only in our body’s motion, but in the motion of external objects (i.e., the ball in our task).

Admittedly, using prism glasses to shift the visual world is not naturalistic. These findings simply represent a step towards more rigorous study of implicit and explicit learning processes “in the wild”. To this end, our study differs from past skill-based learning investigations (11–14) in notable ways. Most importantly, earlier reports (2, 7, 20, 24, 25) have inferred implicit (or procedural) adaptation, under the assumption that the aftereffect during the washout period is entirely driven by implicit processes. This is problematic, as there is no guarantee that participants will stop using explicit strategies, especially when errors are not removed during the washout period which can continue to guide aiming strategies.

Furthermore, there has been no attempt to delineate the potential error sources which could drive subconscious adaptation in a skill-based setting. That is, SPEs and target-errors always co-occur with one another (i.e., the discrepancy between the hand/object and target creates a task error, and a discrepancy between the hand/object and predicted path creates the SPE) and are both known to drive implicit adaptation (1, 3, 7, 10, 15). On the other hand, implicit recalibration may occur through an error-free process such as reinforcement learning. That is, strategies could initially modulate the motor plan, and then be transferred to the implicit system through the development of a use-dependent bias (26, 33). In sum, special experimental conditions must be used to isolate adaptation that can only be attributed to SPEs, which have rarely been applied even in lab-based settings (1, 2, 7).

Here we solved both these issues by adopting the standard best practices in measuring the implicit contribution to adaptation: 1) instructing participants to not use strategy (i.e., aim directly to the target) (3, 8, 30), and 2) removing all visual feedback that could be used to inform them about the success of their movements (34, 35) (in our case, by darkening the room during the no-aiming period). Second, to ensure that this implicit aftereffect was at least in part driven by SPEs, we provided a secondary aiming landmark (2, 7) that helped participants eliminate the target error; adaptation beyond this point must be driven by an SPE, because there is no remaining discrepancy between the ball and the primary target.

The result was a clear implicit aftereffect that exhibited decay properties resembling those studied in lab-based settings. That is, even in the absence of error, the act of moving and the passage of time alone resulted in the partial loss of this memory trial-to-trial (1, 29, 34, 36–38). Further, we could attribute this implicit aftereffect at least in part to SPEs due to the striking overcompensation beyond the primary target (Fig. 2B, overcompensation beyond the target). It is rather remarkable that this distinct SPE-driven phenotype is preserved across motor actions with stark differences in complexity; Mazzoni and Krakauer (2006) studied the movement of an index finger with their arm immobilized via splinting to a tripod, whereas the ball’s path in our task was due to a complicated whole-body coordination that imparted motion on a physical object. In our view, this speaks to the strength and ubiquity of SPE-learning, which we speculate fine tunes our motor behaviors in everyday life.

There are, however, some differences in the magnitude of this aftereffect between our study and that observed by Mazzoni and Krakauer (2). Namely, the 5° implicit aftereffect we detected was smaller than that observed in their task, which reached 20-30°. This dramatic decrease in implicit aftereffect could suggest that in naturalistic tasks, SPE-driven implicit learning is attenuated as the motor skill and corresponding movement complexity increases. In future studies, it will be critical to determine if these aftereffects (roughly 18% of total adaptation) can be increased by changes in experimental factors, or if controlled settings used in laboratory studies promote levels of implicit learning that are not attainable in real-world and skill-based paradigms.

In our task, we suspect that the aftereffects we report in Fig. 2 largely underapproximate the total implicit learning that was achieved during the adaptation period. First, these aftereffects were measured after a substantial break in time (about 1 minute) during which new instructions to aim toward the target were given and the prism goggles were switched to remove the visual shift. Past studies have shown that such breaks in time lead to decay in temporally volatile implicit learning systems (3, 22, 36, 39). Second, the implicit aftereffect was measured after participants had already completed 10 trials without visual feedback, prior to the removal of the shifted prism goggles (see the period between “remove FB” and “goggles off” in Figs. 2A,B). Multiple studies have demonstrated that moving without error feedback also results in a sizeable decay in implicit learning (3, 8, 34, 40). Thus, we strongly suspect that implicit learning is underapproximated by the aftereffect measures in Fig. 2. This is an important limitation of our analysis that could be improved in future work by measuring aftereffects on the very initial no-feedback trials while also minimizing any delay between the adaptation and aftereffect periods.

Apart from these potential bias sources in our aftereffect-based implicit measures, there are critical experimental differences between our work and that of Mazzoni and Krakauer, that could greatly suppress the implicit learning achieved in our ball rolling task. The first factor is temporal delay; given the ∼6.1m displacement between participant and target, the ball arrived approximately 0.75 sec after the motor action had completed. These conditions are known to strongly attenuate implicit learning (8, 20, 21), likely due to the misalignment between motor and error signals in the cerebellar cortex (41–43). Future studies could boost implicit learning by 1) decreasing the distance between the target and participant, 2) requiring faster rolling speeds, 3) or using virtual reality to reduce error latency. The second factor is target error, which is also known to increase implicit learning (1, 3, 10, 44). In Mazzoni and Krakauer (2006), learning occurred slowly without a secondary target (see Fig. 4B, no aiming-target), leading to sustained target errors that were at least 10° or larger throughout the adaptation period. On the other hand, no-aiming-target learning in our task was rapid, reducing the target error to only about 2° within 10 trials or so (Fig. 4A, no-aiming-target). This rapid learning was likely due to strategy use, which was high in our task, about 25° (80% of the total perturbation size). We hypothesize that large explicit strategies deprived the implicit system of target errors between the ball and the task-related target, which would have provided a learning substrate that enhanced the implicit aftereffect. These ideas could be further investigated in future studies by reducing explicit strategies (keeping target errors large) by adding more training targets (45), or limiting movement preparation time (3, 8, 30, 31, 46, 47).

**Figure 4.**
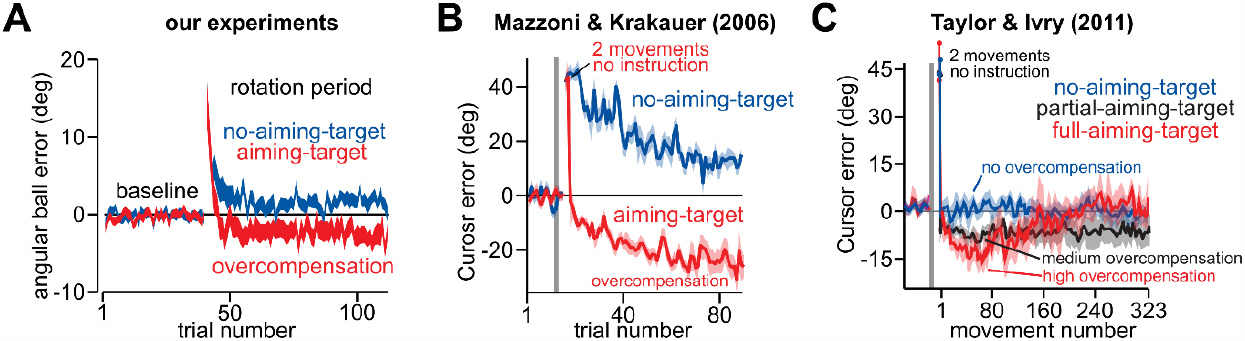
Implicit overcompensation occurs in several experimental contexts. **A**. Trial-by-trial error data from Figure 2A,B with actual target error appearing on the y-axis. Providing an aiming target (in red) caused the angular ball error to deviate beyond the target (i.e., past zero), thus overcompensating for the visual rotation. **B**. Results from Mazzoni and Krakauer (2006), an analogous lab-based experiment, which have been widely cited as evidence for SPE learning. Participants were either given no instruction (blue) or were instructed to use an aiming target to “solve” the rotation and reduce error (red). **C**. Results from Taylor & Ivry (2011), a modified version of the experiment from B. Participants were either given an aiming target during the entire movement (red), an aiming target that disappeared when the movement began (black), or taught how re-aim their movement to counter the rotation without an additional target (blue). Note the graded levels of overcompensation that correspond to the salience of the visual aiming target.

Reduction in implicit adaptation in our experiment could also be inferred based on the time course of overcompensation detected during the aiming-target condition (Fig. 4A, red). Whereas motor behaviors deviated further and further beyond the primary target in Mazzoni and Krakauer (see Fig. 4B, aiming-target), throwing angles in our task (Fig. 4A, aiming-target) remained fixed at nearly a constant 3° overcompensation across the entire adaptation period. The attenuated deviation in our task is entirely consistent with a reduction in the total extent of implicit learning. Indeed, Taylor and Ivry (2011) demonstrated that the extent of overcompensation in an aiming-target condition is directly linked to the extent of implicit learning (Fig. 4C). By providing an aiming target during the entire movement (full-aiming-target), removing it after movement start (partial-aiming-target), or never showing the aiming target during adaptation (no-aiming-target), Taylor and Ivry showed that the extent of overcompensation could be modulated: the more salient the presence of an aiming target, the greater the deviation. These variations in overcompensations were directly linked to changes in implicit aftereffects measured following the adaptation period (not shown in Fig. 4C): the more salient the aiming target during learning, the greater the implicit aftereffect. This again is consistent with past work indicating that SPEs are reinforced by visual target error (1, 3, 10, 44).

Another possible force that attenuates potential overcompensation is a change in explicit strategy. That is, aim reporting protocols have demonstrated late reductions in explicit strategy during training, which presumably functions to prevent the overcompensation that can result from continued SPE-driven implicit learning (24, 25). Similarly, Taylor and Ivry (2011) demonstrated that deviations beyond the target can be eliminated with long exposure to the rotation (Fig. 4C, full-aiming-target). The notion that the explicit system is downregulated over time to counteract SPE-driven implicit learning would be consistent with the order effects we observed in Fig. 3: namely, that initial adaptation rates were greater during the second exposure to the rotation due to savings in explicit strategy, but no change in the terminal residual error could be detected. This hypothesis could be examined in future work by measuring implicit and explicit learning at various points throughout adaptation. Indeed, both our experiment and the other studies illustrated in Fig. 4 provide very limited insight into the time course of implicit learning and explicit strategy, because these are only measured after the rotation period had ended. Measuring the time course of implicit and explicit compensation as they evolve is critically needed to inform our qualitative and quantitative models of SPE-driven adaptation. It is also needed to assess potential limitations in our assumption that implicit and explicit learning are ‘perfectly additive’ processes (22), which may be violated under certain circumstances.

In sum, our work demonstrates that SPE-driven implicit adaptation is not a theoretical construct observable only for constrained motor actions studied in the laboratory. Rather, SPEs are potent drivers of adaptation even during movements that require sophisticated coordination across our entire body as we execute common motor skills. Substantial work remains to better elucidate how SPE-driven implicit learning can be strengthened or attenuated (e.g., via target errors), how implicit processes are coordinated with explicit strategy, and how these two systems evolve over time. Another important future direction will be the application of motion capture to more deeply investigate whether markers of implicit and explicit adaptation can be differentiated across distinct movement components. Initial motion capture efforts have identified a connection between motor learning and whole-body motor adjustments, along with correlations between kinematic variability and learning rates (11). It may be that implicit and explicit processes are differentially expressed across distinct motor control variables (e.g., trunk orientation, ball release timing and variability, wrist and elbow joint angles).

## Acknowledgments

We thank Jenna Floyd, Parker Hill, Erin Kuylenstierna, and Brittney Stone for assistance with data collection. We thank Dr. Shubho Banerjee for helpful data analysis suggestions. Funding provided by a Harrison McCain Emerging Scholar Award (Blustein).

